# Bases for the selection of alternate foot placement during straight- and turning-gait

**DOI:** 10.1101/2024.08.20.608530

**Authors:** Nicholas Kreter, Peter C. Fino

**Affiliations:** University of Utah, Department of Health & Kinesiology, Salt Lake City, UT

**Keywords:** Foot Placement, Stability, Locomotor Control

## Abstract

Humans typically plan foot placement strategy multiple steps in advance when walking across complex terrain. Planning steps in advance is beneficial for both upright stability and forward progress, but one drawback is that new obstacles can make planned foot placement location unsafe between planning and execution, necessitating a rapid shift to foot placement that impacts stability and progress. This study investigates the bases for selection of alternate foot placement during both straight- and turning-gait. Thirteen healthy young adults walked along a virtually projected walkway with precision footholds oriented in either a straight line or with a single 60°, 90°, or 120° turn to the right. On a subset of trials, participants were required to rapidly avoid stepping on select footholds. We found stereotyped alternate foot placement strategies across turn angles that differed by turn strategy. Alternate foot placement for step turns supported the walker’s instantaneous kinematics but not the planned change in trajectory; whereas foot placement for disrupted spin turns supported the walker’s anticipated change in trajectory. We conclude that when humans are forced to rapidly alter previously developed motor plans for foot placement, they utilize a rapid stereotyped behavior that changes based on the demand of the turn.

## 1. Introduction

When walking, humans generally have two goals: (1) the goal of forward progress towards a target destination, and (2) the goal of stability (i.e. stay upright). Both goals are controlled by adjusting foot placement and are thought to be controlled independently. Specifically, forward progress is often viewed as an anticipatory task that can be controlled by changing step length while stability is viewed as an instantaneous task controlled by adjusting step width [1]–[3].

Foot placement is planned several steps in advance when walking through complex environment [4]–[8], which allows humans to utilize visual information to identify obstacles and anticipate a desired path of travel. When obstacles are identified, humans may make subtle changes to anteroposterior (AP) or mediolateral (ML) foot placement to control the direction of instability (i.e., the direction of forward progress) [1], [9]. However, due to the dynamic nature of our world, new obstacles or unrecognized terrain can interfere with planned steps just before foot placement occurs. In such an event, humans must rapidly cancel their initial motor plan, identify a new safe foot placement location, and implement an alternate step – a process referred to here as ‘stepping inhibition’.

Strategies for foot placement during stepping inhibition tasks are stereotyped, with previous research showing a clear preference for alternate foot placement in the sagittal plane. Many stepping inhibition studies interpret the observed foot placement strategy within a constraints hierarchy model where alternate foot placement strategy is influenced by the goals of forward progression, stability, and economy. In such a model, the determinants are ranked relative to task demands, and the strategy used to accomplish the task changes by how the determinants are ranked [10]–[12]. During unobstructed straight-line walking a hypothetical ranking of the determinants might be 1) economy, 2) forward progress, 3) stability. If an obstacle were introduced to the same task, the determinants would be reordered to prioritize stability or forward progress ahead of economy to ensure the obstacle contacted. Using constraints hierarchy models, previous stepping inhibition studies have suggested that a preference for long or short stepping adjustments during a stepping inhibition task are representative of a prioritization of forward progress [10] and a preference for lateral deviations to stepping indicates a prioritization of stability [12]–[14]. However, these conclusions about alternate stepping behavior have been made only when considering straight-line stepping inhibition tasks, where forward progress and stability are assumed to be independent tasks.

When considering gait as a series of falls in the desired direction of travel, one could posit that forward progress is anticipated instability (i.e., the direction the walker anticipates their body will fall to achieve forward progress), and that the goals of forward progress (the direction of instability) and stability (the direction(s) you don’t want to fall in) are directly coupled [15].

Using this definition, perturbations during walking are simply unanticipated shifts to the instantaneous direction of instability that must be corrected. During straight gait, forward progress and stability appear to be independent because typical directions of instability are aligned with anatomical planes of reference. However, humans don’t walk exclusively in straight-lines and often incorporate turns into daily movement. When turning, the direction of instability is allowed to move away from the sagittal plane and into the frontal plane. This process is planned many steps in advance [16], and involves the deceleration of forward momentum, sequential rotation of the head, trunk, and limbs about the stance limb into a new direction of travel, and identification of a safe stepping location for the continuation of forward movement towards a target destination. Generally, humans employ two distinct turning strategies that have different biomechanical demands. Step-turns reorient the body into a new direction of travel while on the outside limb (rotating about the left stance leg for a turn to the right), and spin-turns reorient the body while on the inside limb (rotating about the right stance leg for a turn to the right). Spin turns have been cited as the less stable strategy due to the excursion of the center of mass outside of the base of support [17], [18]. Regardless of strategy, turning impacts temporal gait measures; the amount of time spent in stance phase, the amount of time spent in swing phase, and the swing limb velocity is often asymmetrical between the two limbs when turning [19]. Each of these factors places varying levels of demand on foot placement throughout a turn and may limit the capability of each step to successfully reorient the direction of instability into the desired direction of travel.

When turning, the direction of instability shifts from the sagittal plane towards the frontal plane, with the amount of overlap dependent on the magnitude of the turn. With this overlap, changes to both step length and step width have the ability to influence the direction of instability. As such, strategies for stepping inhibition during a turning task may be different than the stereotyped strategies observed during straight-line stepping inhibition. Here, we investigate alternate foot placement strategies during a stepping inhibition task with turning gait. Specifically, we examine alternate foot placement strategies during both straight- and turning-gait relative to two reference frames: (1) a reference frame aligned with instantaneous state when a stepping inhibition task is identified (stability), and (2) a reference frame aligned with anticipated state (forward progress). This study was exploratory in nature and aimed to investigate (1) the effect of turn angle on alternate foot placement strategy, and (2) whether alternate foot placement strategies differ between step turns and spin turns.

## 2. Materials and Methods

### 2.1. Participants

Thirteen volunteers [7 Female, 13 Right limb dominant, mean (SD) age = 27.2 (3.5) years, mean (SD) mass = 73.8 (15.6) kg, mean (SD) height = 172.8 (7.7) cm] were recruited and screened from the local community. All participants provided written informed consent before participation in this University of Utah IRB-approved study. Inclusion criteria for this study required individuals to be between 18 – 35 years old. Exclusion criteria included (1) a history of neurological pathology that may impact locomotor control, (2) a history of musculoskeletal injuries within the past year that could impact balance, (3) any reconstructive surgery of the lower extremities, (4) and any current medications that could contribute to balance impairments.

### 2.2. Instrumentation and Experimental Setup

A 3D motion capture system (Nexus 2.12, VICON, Oxford, UK) surrounded the walking environment to capture positional marker data at 200 Hz. Participants were outfitted with a custom marker set that featured 41 individual markers and six clusters. Markers were placed on the front of the head (L/R), back of the head (L/R), ears (L/R), acromion process (L/R), lateral epicondyle of the humerus (L/R), forearm (L/R), radial styloid process (L/R), ulnar styloid process (L/R), right shoulder, sternal notch, xiphoid process, C7, T10, posterior superior iliac spine (L/R), iliac crest (L/R), anterior superior iliac spine (L/R), greater trochanter (L/R), lateral epicondyle of the femur (L/R), lateral malleolus (L/R), 1^st^, 2^nd^, and 5^th^ metatarsal heads (L/R), and calcaneus (L/R). Marker clusters were used bilaterally on the shank, thigh, and humerus segments. The marker set was designed such that a full-body 13-segment model could be obtained using the head, upper arms, forearms, trunk, pelvis, thighs, lower legs, and feet. Segments were defined using the normative anthropometric data reported by Dumas et al. [20].

Two projectors (Optoma GT5600; 3600 ANSI lumens, 1080p resolution) projected a custom walking environment on the lab floor with circular stepping targets (radius: 50 mm). A virtually projected square was used to define the origin and global coordinate system to ensure the projectors and motion capture environment were aligned. Positions of the stepping targets were recorded before testing began and checked throughout testing to ensure calibration.

The walking environment was built in Unity (Unity, v2019.4.36f1) and displayed six semi-randomly distributed stepping targets oriented in a straight path (0°), or a path featuring a turn of 60°, 90°, or 120° to the right (Figure 1). The baseline placement of stepping targets was determined by the stepping pattern of the experimenter (NK) walking without visual targets at a brisk pace. To determine the stepping pattern of turn trials, we first calculated the starting position and end position of each turn condition relative to the center of the projected environment. The experimenter (NK) then walked from the starting position to the ending position, turning at the approximate center of the projected turn environment. To scale the walking path to each participants anatomy, step length and width between each target were adjusted as a ratio of the participants leg length (mean (SD) leg length = 911.5 (43.7) mm) divided by the experimenter’s (NK) leg length (996 mm). Leg length was measured as the linear distance between each participant’s ASIS and lateral malleolus of their right leg. Once scaled to leg length, stepping targets one, two, three, and six were randomly jittered within a 150 mm x 150 mm centered on the respective stepping target position and aligned with the walkway. Stepping targets four and five were held to a constant position in the global space for ease of calculating stepping accuracy and strategy in post-processing. For turning trials, turn type was implicit for each trial based on the orientation of the stepping targets. Unity and Vicon Nexus programs were synchronized through a custom code written in C#.

**Figure 1:**
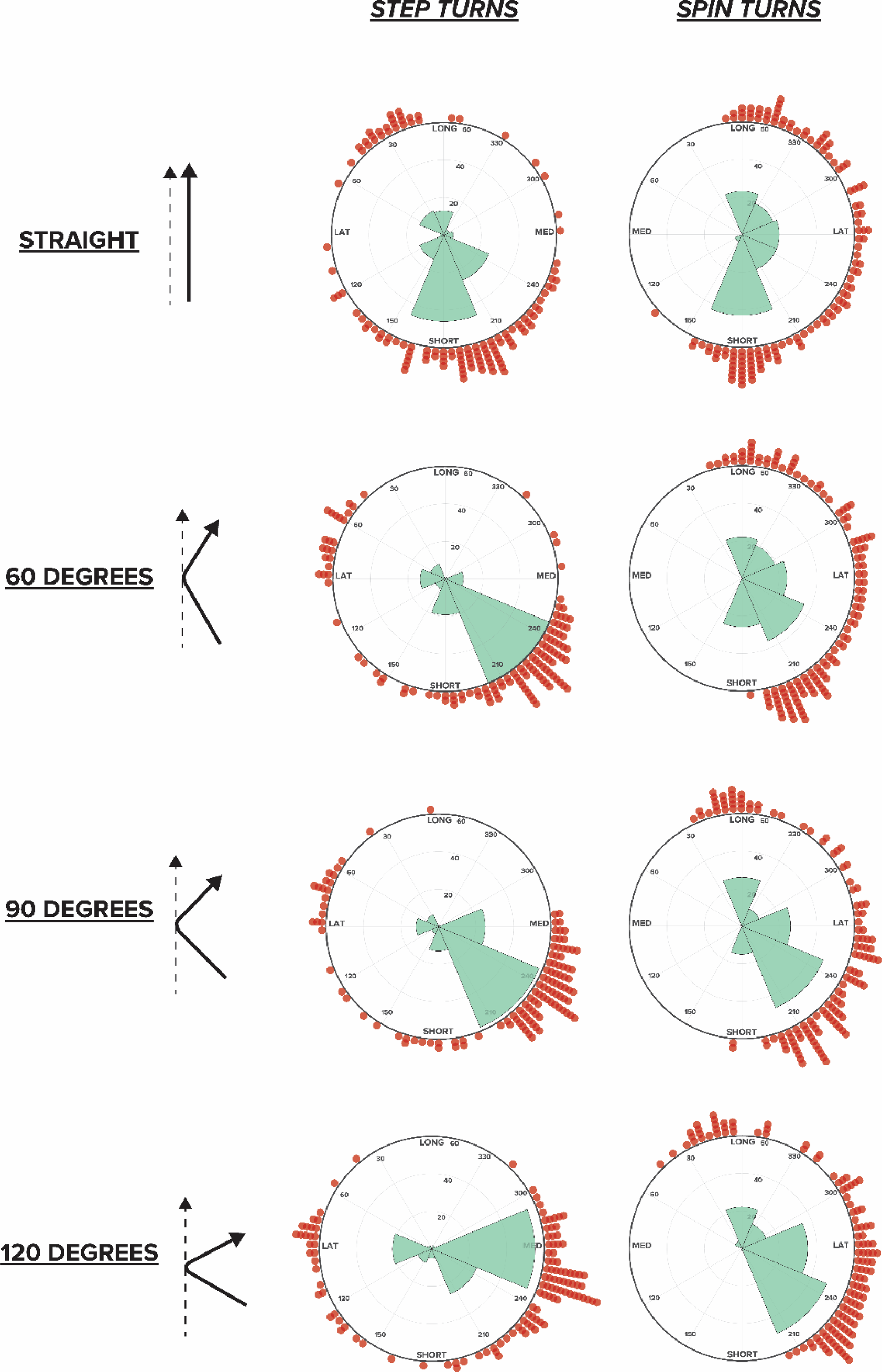
The experimental setup and calculation of alternate foot palcement strategy. Panel A shows example data from an uninhibited walking trial (left) and an inhibited walking trial (right). Grey circles represent precision stepping targets while red circles represent actual foot placement position. For inhibited trials, we calculated the foot placement modification vector (Panel B, top), which is the vector connecting the center of the precision stepping target and the foot placement position. Each foot placement modification vector was then compared against both an approach reference vector (θ_A_) aligned with the direction of travel when the inhibited step was triggered (Panel B, left) or a tangent reference vector (θ_T_) aligned with the anticipated direction of travel for the trial. (Panel B, right). Panel C shows an example trial with a participant walking along the precision stepping targets.

### 2.3. Task and Instructions

Participants were asked to walk along the path of projected stepping targets from a start position to an end position (taped on the lab floor). The starting position was a minimum of two steps from the first precision stepping target in the path, ensuring that gait initiation or starting position had negligible influence on the precision steps of interest. Participants were instructed to walk at a brisk pace, as if “they were trying to catch the bus” [21], to ensure that gait was continuous and each step was not a unique motor event, and also to reduce the possibility of participants slowing to anticipate a perturbation. Participants were also asked to “step on each projected stepping target accurately”, so that the reflective marker on their second metatarsophalangeal joint landed near the center of each stepping target. To ensure that participants were able to walk briskly and accurately, each participant initially completed a series of practice trials where if they stepped such that the marker on their second metatarsophalangeal joint landed within each stepping target the pathway changed color from black to green. If they stepped inaccurately, the course remained black. Practice trials were only performed in a straight-line condition.

Once familiar with walking briskly and accurately, participants were instructed about the remainder of the gait tasks. Participants were told they would be completing walking tasks that required walking along a series of precision stepping targets that were oriented in either a straight path or a path with a turn to the right. Participants were then informed that on a random subset of trials, stepping targets may change color from black to red, and that red stepping targets should be avoided by stepping anywhere else in the environment. Before each trial, participants were instructed to stand in a starting box taped on the floor. Once in the starting box, we projected a path of precision stepping targets for the participant to walk along. Participants started the trial when the experimenter said “begin”. Once participants were standing with both feet in the finishing box at the end of the walkway the trial was ended and participants were instructed to return to the starting box.

### 2.4. Experimental Protocol

Each testing session consisted of 220 trials split evenly between ten blocks. The 220 trials were split so that there were 50 trials per turn angle. Of the 220 trials, 120 were normal, with no steps being inhibited, and 80 trials with a stepping inhibition task on the fourth step (the step in the middle of the turn for turning trials). The remaining 20 trials were “catch” trials where steps either before or after the turn were inhibited, to prevent participants from solely focusing on the middle step of each trial. Half of the trials were displayed by the projector as step turns and the other half were displayed as spin turns. Inhibited step turns only disrupted the left foot and inhibited spin turns only disrupted the right foot. Blocks were organized so that there were five trials of each turn angle (three normal trials and two inhibited trials), and two “catch” trials in each block. Each within-block trial order was randomized to make the task less predictable. The catch trials were controlled so that the first catch trial of each block would occur randomly between trials 4-7 and the second would occur between trials 15-18. Inhibited steps changed color from black to red when the reflective marker on the second metatarsal head entered within a one-step radius of the step to be inhibited (See Supplementary Video), which coincided with foot contact on the n-1 step.

### 2.5. Processing

All marker data was processed in Vicon Nexus (v.2.12). Filtering, gait events, and outcome measures were computed with a custom code in Matlab (v. R2020b, The MathWorks Inc., Natick MA, USA). Kinematic data were filtered using a 4^th^ order low-pass Butterworth filter with a cutoff frequency of 6Hz. Heel strike and toe-off events were determined with a maximal displacement algorithm adapted for turning gait [22], [23]. To determine these gait events during turning trials, positional marker data were rotated to align with the whole-body center of mass velocity.

For inhibited steps, the vector connecting the center of the original stepping target and the position of the marker on the second metatarsal head at mid-stance defined a foot placement modification vector (FPMV). We established two walkway-based vectors. One vector connected the starting position of the trial and the apex of the turn and defined the approach reference frame (Figure 1B; aligned with the walker’s immediate state when the perturbation is presented). The other vector was oriented tangent to the turn angle and defined the tangent reference frame (Figure 1B; aligned with the goal of forward progress when the perturbation is presented). For straight-gait trials these reference frames are the same and aligned parallel to the walkway. The angle between the FPMV and each walkway vector was calculated and called a foot placement modification angle (FPMA). Each FPMA was then binned into one of eight potential foot placement strategies, long, short, medial, lateral, long-lateral, long-medial, short-lateral, and short-medial.

### 2.6. Statistical Analysis

Descriptive and inferential statistics were calculated using the circular statistics toolbox in Matlab [24]. Circular mean and mode were reported as measures of central tendency (Table 1). For each turn angle and turn type, variance is reported as the circular standard deviation (Table 1). To our knowledge, there are no frequentist hypothesis tests that appropriately compare the means of repeated measures periodic (i.e., circular) data. Therefore, we used a hierarchical bootstrapping method, described by Saravanan and colleagues, to compute circular means and 95% confidence intervals of FPMA for each condition (10,000 samples with replacement) [25], [26]. The hierarchical bootstrap first samples with replacement from the subject level. Once the subject is selected, that subject’s trials are sampled with replacement for the same number of trials as was performed in the experiment. Since there are no hypothesis tests that can handle repeated measures periodic data, we instead report descriptive statistics from our bootstrapping analysis, and also performed circular correlation analyses to assess the association between turn angle and mean FPMA for step turns and spin turns in both reference frames.

**Table 1:** Circular mean, standard deviation, and mode FPMA for step turns and spin turns in each reference frame.

|  | Approach Reference Frame |  |  |  | Tangent Reference Frame |  |  |  |
| --- | --- | --- | --- | --- | --- | --- | --- | --- |
|  | Step Turns |  | Spin Turns |  | Step Turns |  | Spin Turns |  |
| **Condition** | Mean (SD) | Mode | Mean (SD) | Mode | Mean (SD) | Mode | Mean (SD) | Mode |
| Straight (0°) | 199° (74°) | 196° | 270° (68°) | 180° | 195° (77°) | 196° | 277° (67°) | 180° |
| 60° | 178° (57°) | 188° | 237° (57°) | 164° | 206° (63°) | 228° | 274° (57°) | 196° |
| 90° | 179° (56°) | 196° | 227° (60°) | 164° | 226° (62°) | 236° | 280° (61°) | 212° |
| 120° | 177° (64°) | 188° | 217° (51°) | 164° | 235° (68°) | 252° | 283° (52°) | 220° |

## 3. Results

Mode FPMA revealed preferred foot placement strategies for all turn angles and turn types for both the approach and tangent reference frame definitions (Table 1). The mode FPMA for step turn in the approach reference frame were 196° for straight trials, 188° for 60° turns, 196° for 90° turns, and 188° for 120° turns. For spin turns in the approach reference frame, the mode FPMA was 180° for straight gait, and 164° for each of the other turn angles. The mode FPMA for step turns in the tangent reference frame were 196° for straight trials, 228° for 60° turns, 236° for 90° turns, and 252° for 120° turns. For spin turns in the tangent reference frame, the mode FPMA was 180° for straight gait, and 196° for 60° turns, 212° for 90° turns, and 220° for 120° turns.

The hierarchical bootstrapped circular mean FPMA for step turns in the approach reference frame were 181.9° for straight gait, 175.3° for 60° turns, 176.8° for 90° turns, and 175.1° for 120° turns (Table 2). For spin turns in the approach reference frame circular means from the hierarchical bootstrap analysis were 244.3° for straight gait, 226.7° for 60° turns, 210.2° for 90° turns, and 206.3° for 120° turns. For step turns observed in the tangent reference frame, the circular mean FPMA was 181.9° for straight gait, 205.5° for 60° turns, 221.6° for 90° turns, and 235.4° for 120° turns. For spin turns in the tangent reference frame we observed a circular mean FPMA of 244.3° for straight gait, 256.5° for 60° spin turns, 255.8° for 90° spin turns, and 266.3° for 120° spin turns.

**Table 2:**
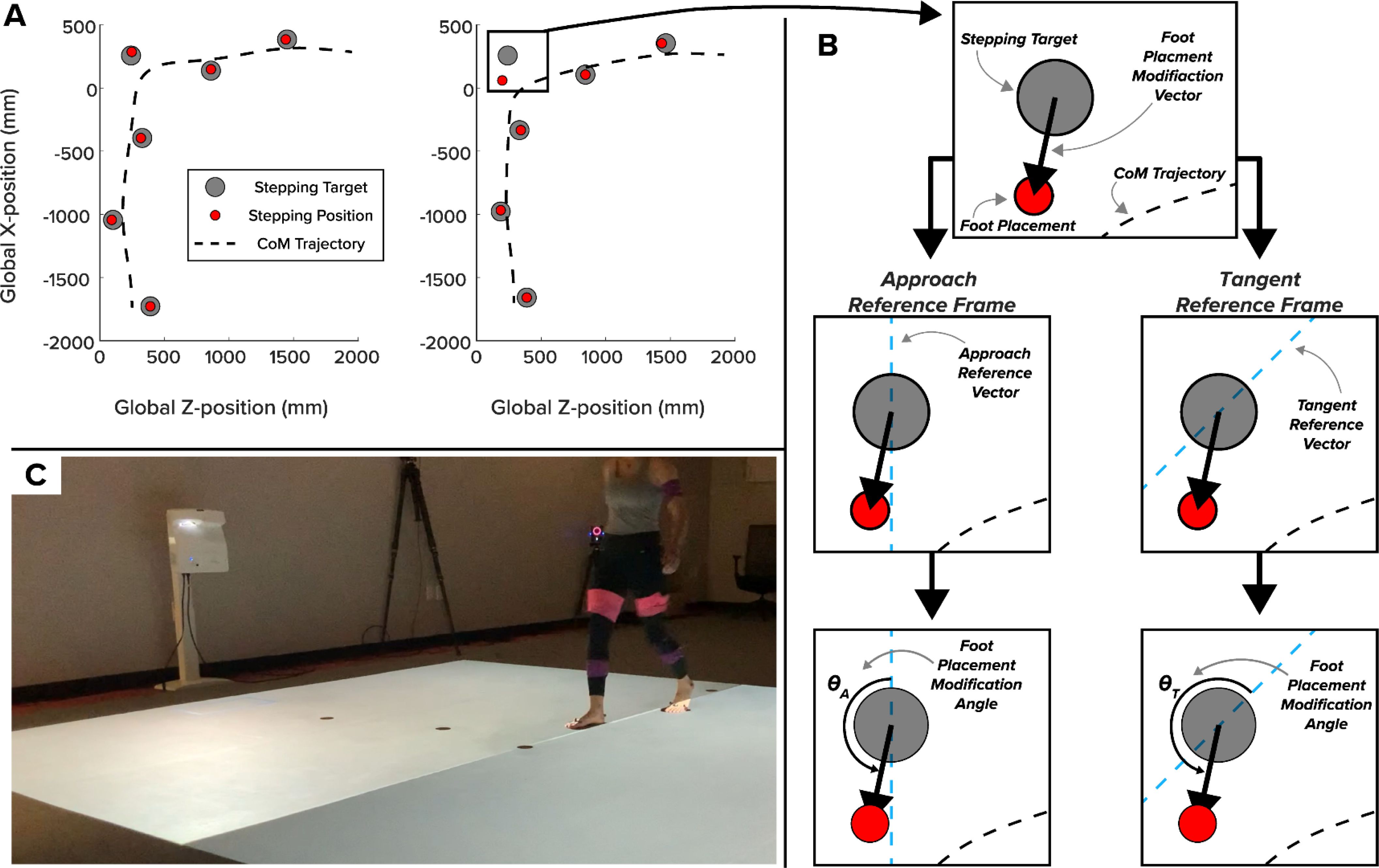
Circular mean and 95% confidence intervals of the hierarchical bootstrapped means for each turn type and reference frame.

For step turns in the tangent reference frame, we observed a consistent change in the circular mean strategy as turn angle increased. For each turn angle in this reference frame, the shift of the mean FPMA was similar to the rotation of the reference frame relative to the straight condition. Relative to the straight gait condition, we observed a shift of 23.6° to the FPMA for the 60° turn (tangent = 30° from straight), 39.7° for the 90° turn (tangent = 45° from straight), and 53.5° for the 120° turn condition (tangent = 60° from straight). For spin turns in the approach reference frame we observed a similar, albeit less consistent, shift in mean FPMA: 17.6° for the 60° turns, 34.1° for the 90° turns, and 38° for the 120° turn conditions.

Circular correlation analyses of bootstrapped means showed that turn angle explained the greatest proportion of variance for inhibited step turns where FPMA was calculated in the tangent reference frame (r^2^ = 0.60) and inhibited spin turns in the approach reference frame (r^2^ = 0.40). Turn angle explained the least variance for inhibited step turns when FPMA was calculated in the approach reference frame (r^2^ = 0.01) and inhibited spin turns in the tangent reference frame (r^2^ = 0.13).

## 4. Discussion

The objective of this exploratory study was to investigate the effect of turn angle and turn type on alternate foot placement strategy following an inhibited step during straight and turning gait. Generally, we found stereotyped foot placement strategies regardless of turn angle for both step turns and spin turns. Mode strategy showed a strong preference for short steps during inhibited step turns and short-lateral steps for inhibited spin turns (Figure 2 and Figure 3). Stepping strategy for spin turns was more varied than for step turns, but still showed similar behavior (Figure 2 and Figure 5). While the strategies for step turns and spin turns appeared similar at face value, hierarchical bootstrap analyses revealed differences between step turns and spin turns that suggest they are planned and controlled in fundamentally different ways (Figure 4). Specifically, step turns appear to be controlled relative to a walker’s instantaneous CoM state while spin turns appear to be controlled relative to a walker’s anticipated CoM state.

**Figure 2:**
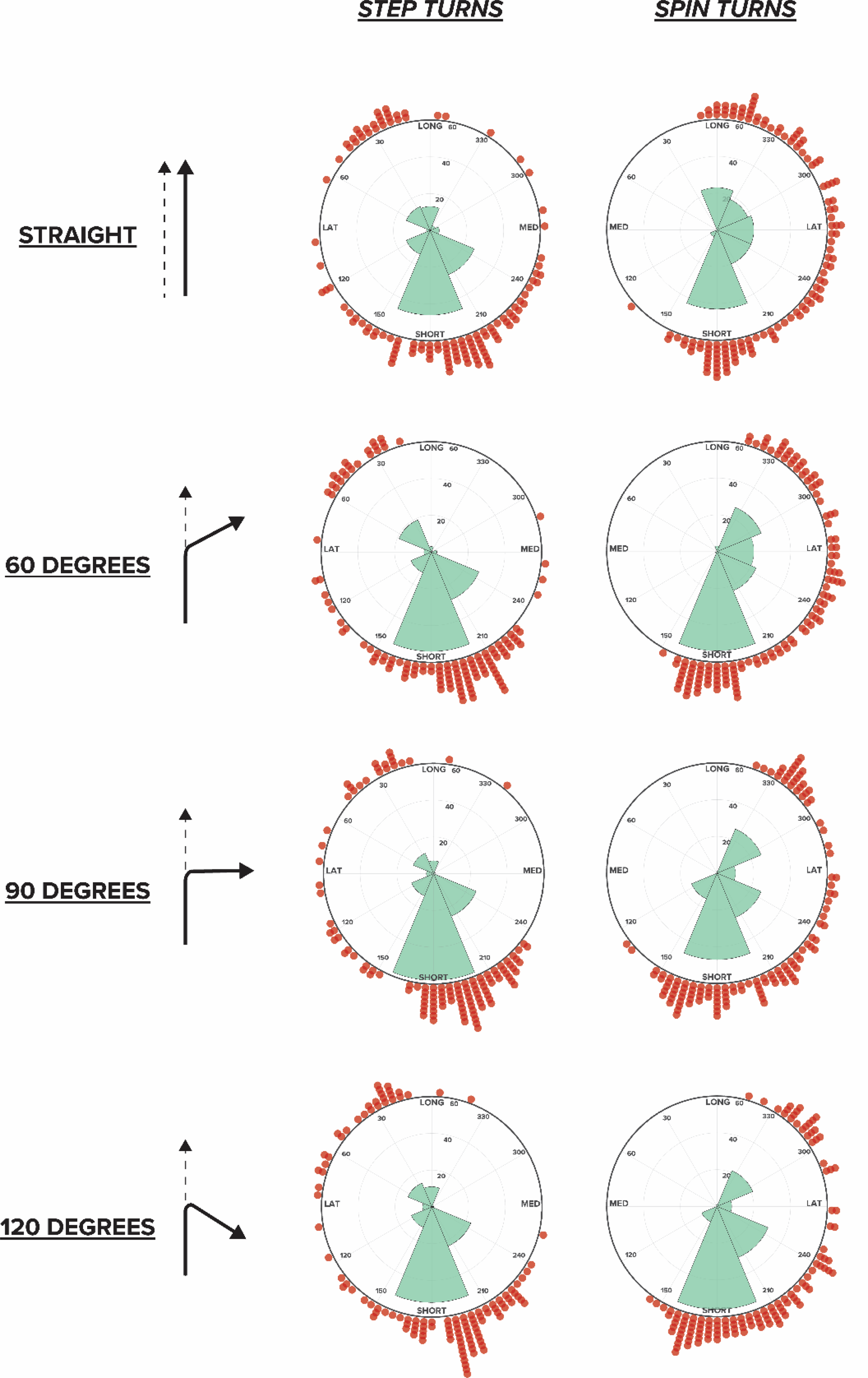
Distribution of alternate steps in the tangent reference frame. Turn angles are shown by turn angle and turn type. Polar histograms (green) present the alternate stepping strategy distribution for the eight categorical bins. Individual trials (red) show the distribution of alternate stepping strategy in 4° bins. For straight trials “step turns” inhibited the left foot and “spin turns” inhibited the right foot.

**Figure 3:**
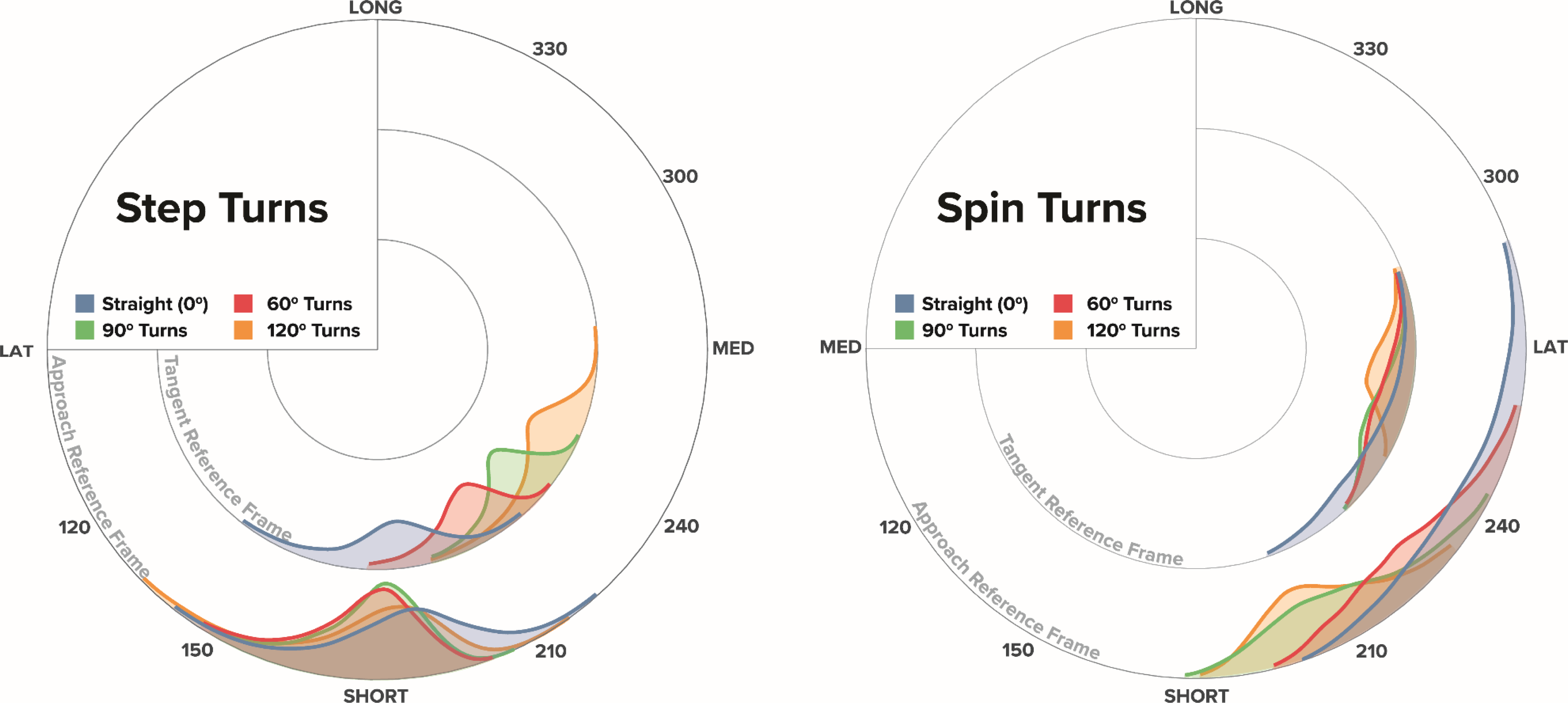
Distribution of alternate steps in the approach reference frame. Turn angles are shown by turn angle and turn type. Polar histograms (green) present the alternate stepping strategy distribution for the eight categorical bins. Individual trials (red) show the distribution of alternate stepping strategy in 4° bins. For straight trials “step turns” inhibited the left foot and “spin turns” inhibited the right foot.

**Figure 4:**
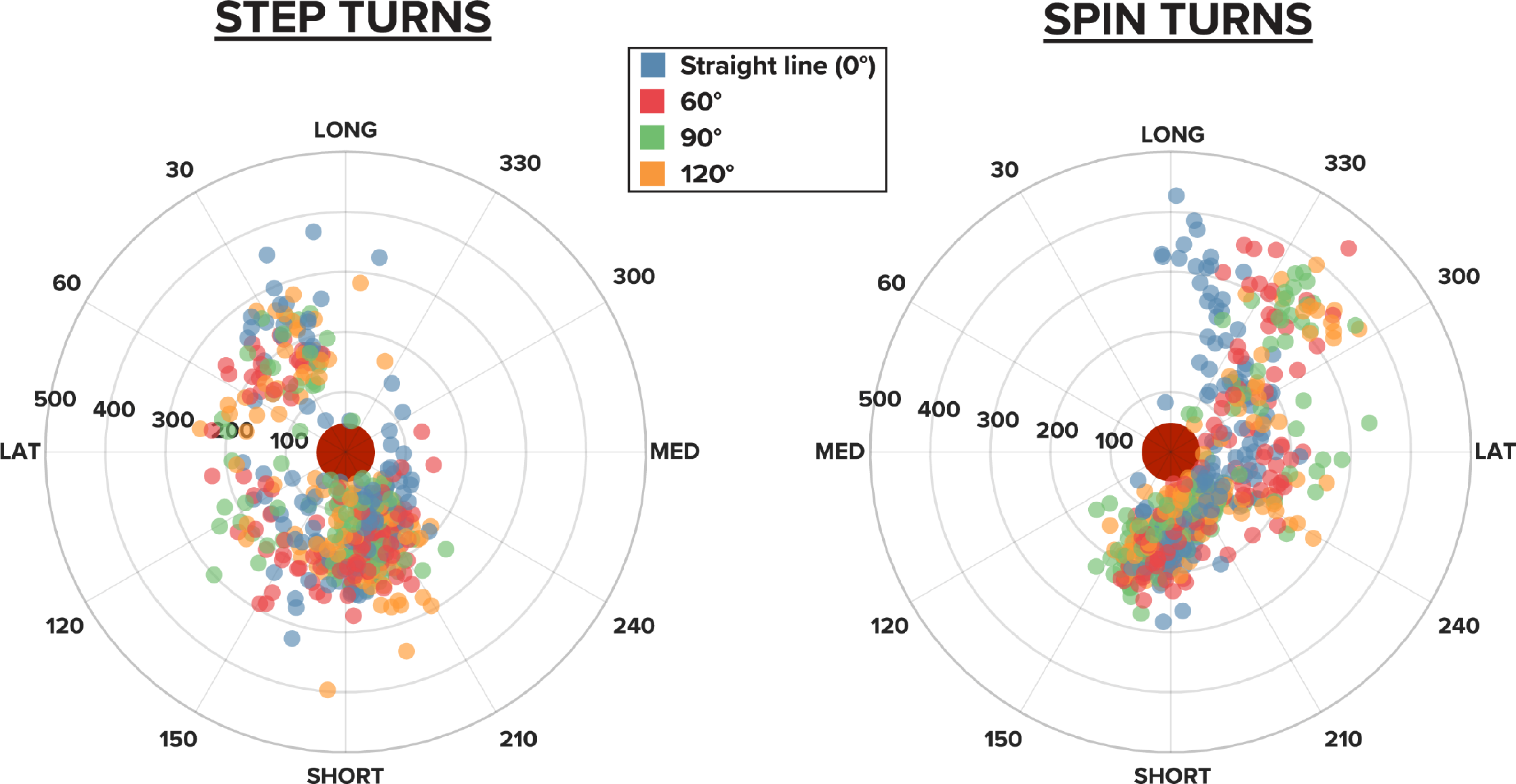
95% confidence intervals of hierarchical bootstrapped means for each reference frame and turn type. Data are presented for straight gait (blue), 60° turns (red), 90° turns (green), and 120° turns (orange) for both step turns (left) and spin turns (right). The inner ring of each circle represents the means and confidence intervals for data examined in the tangent reference frame while the outer ring represents data from the approach reference frame. Extreme edges of each distribution represent the upper and lower bounds of the 95% confidence intervals for each distribution.

**Figure 5:**
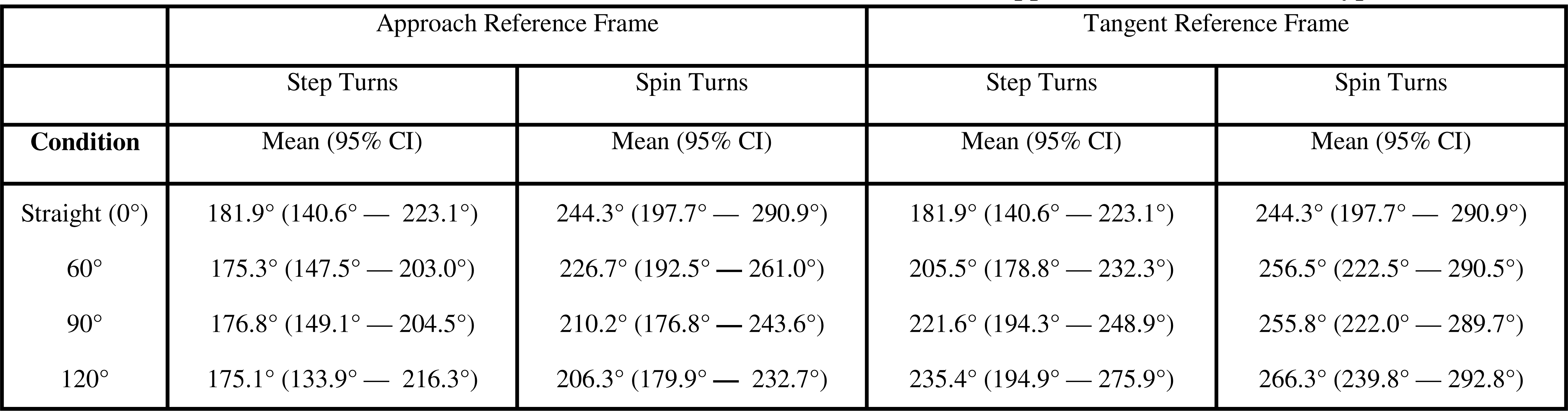
Location and magnitude of all inhibited steps in the approach reference frame. Steps from straight (blue), 60° (red), 90° (green), and 120° (yellow) turning trials are presented relative to the inhibited stepping target (central red dot). Each ring in the polar plot represents a 100 mm change from the center of the stepping target. Each spoke of the polar plot represnts a 30° change around the arc. All data are presented from the approach reference frame.

### 4.1. Alternate Foot Placement Strategy During Turning

When assessed in the approach reference frame, our results generally agreed with Patla’s original model for alternate foot placement behavior [10]. Here, the participants’ dominant choice for alternate foot placement were in-line with the direction of travel when the inhibited step was triggered, with a preference for short steps over long steps. Long and short adjustments are quicker adjustments to implement for inhibited steps because they take advantage of muscles that are already active during the swing phase of straight gait; primarily the flexors and extensors of the knee and hip [27]. Medial and lateral adjustments require use of the hip adductors and abductors, respectively, muscles not commonly active during swing phase of gait [2], [27].

The preference for short steps in this study is a slight break from Patla’s original model [10]. Long steps are usually the preferred choice during inhibited stepping tasks [10], [12], [28], [29] because they allow participants more time to plan foot placement and reduce burden on the execution of future steps. Short steps are typically less favorable because they run the risk of excessive forward angular momentum if forward linear momentum isn’t reduced before foot placement. However, such conclusions on the pros and cons of long and short steps have only been discussed with regard to straight gait. During turns, CoM dynamics are different and short steps may be more favorable [30]. First, turning is planned many steps in advance [16] and necessitates a reduction of sagittal-plane momentum to reorient the direction of instability [31]. A detriment of a short stepping strategy during straight-gait is the risk of excessive angular momentum in the sagittal plane, but during a turn, that momentum is already being reoriented away from the sagittal plane to a new direction. Since short steps don’t hinder forward momentum or create unexpected sharp changes to trajectory, they may also be less costly from an energetic perspective [32]. An additional benefit of a short step during a turn is that it provides a walker with immediate influence over one’s kinematic state, whereas long steps delay a potentially necessary change to CoM-trajectory during turning. Since the step that was inhibited in this experiment was the step responsible for reorienting body trajectory, a short step strategy may allow the initiation of the reorientation of body state sooner and reduce the necessity of the recovery step to reorient the body and maintain upright posture.

While our results generally agreed with the results reported by Patla and Moraes [14], our overall conclusions differ slightly. Patla [10] and Moraes [11]–[13] originally interpreted their data with the assumption that forward progress and stability are independently controlled by foot placement. That is, long or short adjustments are related to forward progress, while medial and lateral adjustments are related to stability. This assumption is only valid for straight-gait tasks and is not compatible with turning-gait because body trajectory and body orientation don’t necessarily align. Therefore, determining the influence that a change to foot placement has on forward progress or stability is quite difficult and can change with different reference frames [33]. When defining forward progress as anticipated instability straight- and turning-gait tasks can be interpreted within the same framework. When the two goals are directly coupled, individuals are allowing instability tangent to the path of travel and preventing instability in all other directions. In this definition, neither forward progress nor stability can be prioritized, and instead conclusions must focus on whether stepping behavior is influenced by anticipation of future CoM state, or influenced by CoM state when the obstacle is identified. In the present study, we found that the influence of instantaneous CoM state and anticipated CoM state differed depending on what style of turn was executed.

### 4.2. Differences Between Step Turns and Spin Turns

Alternate foot placement appeared to be controlled differently between step turns and spin turns. Hierarchically bootstrapped circular means revealed a consistent shift in FPMA as turn angle increased in the tangent reference frame for step turns and the approach reference frame for spin turns. While these shifts in mean FPMA may look like a shift in functional stepping strategy, the shift in FPMA for each turn angle was similar to the angle of a vector tangent to the turn (i.e., 30° for the 60° turn, 45° for the 90° turn, and 60° for the 120° turn) and was likely driven by the rotation of the reference frame rather than an actual change in stepping strategy. Counterintuitively, the reference frames in which we see no change to alternate foot placement across turn angles likely reflect the reference frame in which walkers decided their alternate foot placement strategy. For step turns we observed no change in strategy in the approach reference frame, indicating a strategy that controls instability in the walker’s instantaneous state when encountering the disrupted step. Correlation analyses further supported this conclusion, showing that turn angle explained 60% of the variance in alternate foot placement for step turns in the tangent reference frame, but only 1.5% of the variance for step turns in the approach reference frame. Conversely, spin turns showed minimal change to mean strategy in the tangent reference frame, suggesting a control mechanism aimed at preserving the anticipated change to instability throughout the turn.

The observed difference between step and spin turns may indicate fundamentally unique control strategies for perturbations during step turns and spin turns. This explanation pairs well with previous work exploring the kinematic differences between the turn types [17], [30], [34]. Due to the compromised position of the CoM and contralateral limb during spin turns, walkers experiencing a sudden change to foot placement (like an inhibited step) have little ability or opportunity to correct a spin turn once it has been initiated [17], [30]. Alternate stepping strategies that align with anticipated changes to CoM state may disrupt frontal- and/or sagittal-plane kinematics but should preserve the change in direction of travel. Since there is little a walker can do during a spin turn to correct for a perturbation, continuing along their planned direction of travel until the next opportunity to step may be safest. The relationship between the CoM and BoS during step turns allows the walker to generate torque throughout stance phase and immediately correct for any perturbations that arise [8], [17]. During a step turn with an inhibited step, an adjustment to foot placement that aligns with instantaneous CoM state may come with an immediate cost to the planned change of direction but ensures frontal- and sagittal-plane dynamics remain unchanged. Kinematic analyses in our companion manuscript largely support these interpretations. Briefly, we found long lasting disruptions to frontal and sagittal angular momentum, but no change in the transverse plane, following inhibited spin turns [35]. For step turns we observed an immediate but brief disruption to transverse angular momentum, but little-to-no impact on frontal and sagittal dynamics. We also observed a greater frequency and magnitude of stepping error on the first recovery step after inhibited spin turns, suggesting that the disruption caused by the inhibited step wasn’t corrected until after the turn.

Due to the excess threat to stability during spin turns, we expected that the shift to foot placement strategy would be opposite between spin turns and step turns, with step turns shifting medially and spin turns shifting laterally. During step turns the majority of alternate steps were adjusted in the AP direction, with a heavy preference for short steps. While we also observed a preference for short and short-lateral steps during spin turn maneuvers, the distribution of alternate steps showed a greater frequency of lateral foot placement adjustments than during step turns. One explanation for the different distributions of alternate steps between step and spin turns is due to differences in threats to stability; lateral steps during a spin turn may help reclaim some stability. However, the difference in foot placement distribution could also be a result of limb dominance or lateralized limb control during walking.

The distribution of alternate steps along the lateral edge of the stepping target for spin turns may also be due to lateralized control of gait. Lateralized use of the lower extremities, where one limb prioritizes stabilization and the other prioritizes control or mobilization, has been proposed across many gait studies, and is often tied to limb dominance [36], [37]. During turning maneuvers, lateralized control of the limbs is necessary to change direction [38], [39]. If lateralization during gait is tied to limb dominance, changes to foot placement of the stabilizing limb may appear different to changes of the mobilizing limb (i.e., the limb predominantly responsible for propulsion of the CoM) during an inhibited stepping task. Specifically, the mobilizing limb may exhibit more precise control tailored to instantaneous CoM dynamics. In the present study, a change in foot placement was necessary for both the left limb (step turns) and the right limb (spin turns), and foot placement strategy for spin turns appears unimodal for each of the turn angles whereas the foot placement strategy for step turns appears bimodal. The greater frequency of stepping adjustments in the ML direction for spin turns may indicate that the dominant limb is weighted towards finer motor function tied to mobilization or stability while the non-dominant limb is heavily weighted to control gross function like propulsion or forward progress [36], [37], [40].

### 4.3. Limitations

This study should be interpreted within the context of its limitations. First, instructions on walking speed and how to step accurately may have influenced decisions for alternate foot placement. We asked participants to walk at a brisk pace and step so that a marker on their second metatarsal head would be placed in the center of the stepping target. The brisk pace was intended to ensure participants didn’t slow in anticipation of inhibited steps but may have also influenced kinematics to the point that a different alternate stepping strategy was preferable relative to a comfortable pace. Further, the instructions to step accurately may have induced a preference for the short step by adjusting the minimal displacement of the foot. To this point, it is unclear if the minimal displacement criteria for alternate steps is controlled relative to the center of mass of the foot, or a different anatomical landmark, like the ball of the foot or heel, that is more directly involved in the process of stepping. For participants with larger feet, the marker for stepping accurately was likely a few centimeters anterior to the center of mass of the foot. Second, this project relied heavily on the synchronization of many pieces of technology. The communication between Vicon and Unity was likely not instantaneous, and small changes in the latency of the inhibited step trigger were not recorded. While likely occurring within a sub-perceptible threshold, this could have led to changes in the available response time for inhibited steps on some trials.

## 5. Conclusion

We found that when walkers need to rapidly avoid an obstacle in a planned stepping location, they regularly execute a stereotyped foot placement strategy. In contrast with previous studies that suggested forward progress or stability are prioritized dependent on task constraints, our results suggest that humans don’t necessarily prioritize stability or forward progress, but instead determine adjustments to foot placement based on turn type. Specifically, we observed related strategies between step turns and spin turns; however, alternate foot placement strategies for step turns were selected to control instantaneous CoM state while strategies for spin turns were selected to control the anticipated CoM state. Our findings suggest that these turn types contain fundamental differences in how they are planned and actively controlled in the presence of perturbations.

## Supporting information

Supplementary video showing an inhibited stepping trial in slow motion

## 6. Acknowledgments

The authors would like to acknowledge the contributions of Paula Kramer and Cameron Jensen for their assistance with data collection throughout this project. We also extend our thanks to Nora Fino for providing biostatistical consultation.

